# Spatial heterogeneity in resources alters selective dynamics in *Drosophila melanogaster*

**DOI:** 10.1101/2020.09.05.283705

**Authors:** Audrey E Wilson, Ali Siddiqui, Dr. Ian Dworkin

## Abstract

Environmental features can alter the behaviours and phenotypes of organisms and populations evolving within them including the dynamics between natural and sexual selection. Experimental environmental manipulation, particularly when conducted in experiments where the dynamics of the purging of deleterious alleles are compared, has demonstrated both direct and indirect effects on the strength and direction of selection. However, many of these experiments are conducted with fairly simplistic environments when it is not always clear how or why particular forms of spatial heterogeneity may influence behaviour or selection. Using *Drosophila melanogaster*, we tested three different spatial environments designed to determine if spatial constraint of critical resources influences the efficiency of natural and sexual selection. We conducted two allele purging experiments to 1) assess the effects of these spatial treatments on the selective dynamics of six recessive mutations, and 2) determine how these dynamics changed when sexual selection was relaxed and the spatial area was reduced. We found that allele purging dynamics depended on spatial environment, however the patterns of purging rates between the environments differed across distinct deleterious mutations. We also found that for two of the mutations, the addition of sexual selection increased the purging rate.

## Introduction

Understanding mating systems and the dynamics between the sexes can illuminate how sexual selection acts within populations, driving many organisms’ behaviours and phenotypes. Key work in the theory of mating systems conducted by Bateman (1948), Trivers (1972), and Emlen and Oring (1977) has led many studies being dedicated to examining male and female interactions across different species and populations. The mating systems of numerous species have been shown to vary due to local adaption or ecological constraints due to environmental factors (Miller and Svensson 2014). For example, ungulate species that inhabit open environments tend towards group mating systems while those within closed or forested environments tend to adopt small group or pair mating systems (Carranza 2000; Bowyer et al. 2020). This variation in behaviour can occur within species as well, as seen in the mating system of *Prunella modularis,* which has been shown to shift between polygyny, polygynandry, and polyandry depending on food distribution (Davies and Lundberg 1984). Environmental features such as spatial size, structure, resource abundance, and climate can alter the strength of sexual selection and conflict (both intra- and inter-) on an individual, in turn leading to fitness payoffs for certain phenotypes. In *Sancassania berlesei*, increasing environmental complexity changes the fitness differences between the fighter and scramble male morphs which was believed to be a result of reduced encounters between fighter males (Lukasik et al. 2006). Another example can be found in certain populations of katydid, where sex role reversal occurs under conditions of low resource abundance, placing a greater influence of inter- and intra-sexual selection on females (Gwynne and Simmons 1990). Since environmental variation can impact fitness, it is important to keep environmental context in mind when studying the strength of natural selection, sexual selection and sexual conflict.

Along with the environment, understanding the interaction between sexual selection and other components of natural selection (fecundity and viability) is important for determining an organisms’ or a populations’ phenotypic and behavioural origins. Since the term was introduced by Darwin (1871), studies have focused on how traits under strong sexual selection (weaponry, ornaments, and mating behaviours) arise and persist within populations. When sexual conflict is present, mutations may be beneficial in one sex but deleterious in the other (antagonistic pleiotropy), allowing for the maintenance of conditionally deleterious alleles. In many species, an extreme case of this is males harming females during copulation, either through mating itself or ejaculates, in order to prevent re-mating, further securing the males’ paternity (Johnstone and Keller 2000). However, while often portrayed as being at odds with one another, individuals of higher overall condition will on average receive more mates, resulting in sexual selection working in tandem with other components of natural selection. For instance in ungulates, males of overall higher condition tend to have the largest weaponry and are better able to obtain fertilizations along with access to females themselves (Preston et al. 2003; Hoem et al. 2007; Vanpé et al. 2007; Emlen 2008).

A common way of determining how various factors influence natural and sexual is to conduct allele purging experiments. Within these experiments, deleterious mutations are introduced into populations at a known frequency (or via induced mutations) and the rate they are removed from the populations over time is recorded or populations undergo various fitness assays. Experimental conditions are manipulated (thermal stress, dietary stress, population density, environmental complexity, and mate choice (Sharp and Agrawal 2008; Wang et al. 2009; Young et al. 2009; Hollis et al. 2009; MacLellan et al. 2009; Laffafian et al. 2010; Hollis and Houle 2011; McGuigan et al. 2011; Arbuthnott and Rundle 2012; Clark et al. 2012; Maclellan et al. 2012; Singh et al. 2017; Colpitts et al. 2017) and purging rates (or fitness) are compared to obtain estimates of the effects these conditions have on selective dynamics. While several kinds of these studies have been conducted, many show contrasting results in reference to whether sexual selection aids natural selection in the removal of deleterious alleles. One potential reason for such inconsistencies is that most experiments are performed in small, simple environments (i.e. small vials) at relatively high densities, and it is not clear the degree to which this may influence the strength and orientation of selection. Such simple and high-density environments likely constrain individuals in terms of mating strategies available in more natural conditions. Alternative mating strategies are density-dependent in several species (Greenfield and Shelly 1985; Höglund and Robertson 1988; Kokko and Rankin 2006), and particularly for *Drosophila melanogaster*, territorial defence strategies by males are less likely to occur when the population is at a high density (Hoffmann and Cacoyianni 1990). Simple environments may also influence female strategies in that they may accept more mates due to being unable to seek refuge or escape from constant male harassment (Byrne et al. 2008). Creating a more “complex” environment consisting of a larger space, multiple food cups, and additional spatial structure to alter the interactions between the sexes, Yun et al. (2017) showed that female harassment of high quality *D. melanogaster* females was greater in the simple fly vial environments used in many experiments, exaggerating the effects of sexual selection to reduce variance in female fitness.

Since Yun et al.’s (2017) experiment, there have been several studies conducted to determine how natural and sexual selection changes within simple (high density in single vials or bottles) versus “complex environments” (lower density cages with multiple resources for interactions to occur). In a later study, Yun et al. (2018) found flies that had mating opportunities within “complex” environments adapted more quickly to novel larval environments as opposed to those mating in simple environments or lacking mate competition. Using a similar environmental design but creating a larger, lower density simple environment, Colpitts et al. (2017) demonstrated that “complex” environments aided the purging of two deleterious mutations that had previously been found to have no difference in purging rate while manipulating opportunity for mate choice (Arbuthnott and Rundle 2012). Singh et al. (2017) showed increased purging rate of deleterious alleles from populations evolving within these “complex” environments, while MacPherson et al. (2018) revealed that low quality females experienced a greater reduction in fitness due to male harm compared to high quality females but only in “complex” relative to simple environments. These studies exemplify that with even modest changes in spatial environment (increasing space and lowering density of individuals), the dynamics of natural and sexual selection can vary vastly. Complexity without the manipulation of overall environment size has been shown to influence female fitness in terms of offspring production (Malek and Long 2019), but this has not been used to test overall population fitness. While these studies potentially show how these forces interact in a way that may be more representative of what is seen in nature, the types of environments employed are still simple and largely reflect changes in density. However, it is important for such experiments to explicitly consider factors that are known to influence mating strategy as well, such as territory availability and spatial heterogeneity of resources.

Increasing the environmental complexity in which populations evolve may reveal new patterns of how sexual selection acts, particularly for *D. melanogaster*, which as a species shows considerable variability in mating strategy in different spatial contexts. Typically displaying scramble competition in the lab, territorial behaviours and resource defense polygyny have been observed when *D. melanogaster* males are given a desirable resource (Hoffmann 1987). Males also appear to display this behaviour more often when females are present, when there is a low density of males, and the resource is readily used by females for oviposition and resource patches are in a range of sizes (~20mm diameter) (Hoffmann and Cacoyianni 1990). Within laboratory experiments, where aggressive interactions amongst *D. melanogaster* males are observed, it is typical that larger males or males that hold residence of a territory first, have greater reproductive success (Hoffmann 1987). Considering this, if populations are within an environment that allows males to benefit from territorial behaviour, these populations may show an increase in overall fitness, and more variation in mating strategies. Yet to date, most experimental evolution and purging experiments have not considered these explicit factors in their design.

While the previous work outlined above has made considerable contributions to our understanding of the interplay between environmental complexity and selective forces, the environments used in these experiments are relatively simplistic when considering the plasticity of animal mating behaviour. We conducted a series of short term allele purging experimental evolution assays where environmental complexity and the accessibility of *D. melanogaster* to critical resources were manipulated with these factors in mind. In the first part of this experiment we looked at how differences in resource patch size and accessibility influenced the purging of six recessive deleterious mutations from populations being held within a series of complex environments. Specifically, we provided multiple resource patches of high (to maximize female fecundity) and low quality. In each treatment high quality patch size and accessibility varied according to how they should potentially influence aspects of territoriality. In the second experiment, we examined how the rate of removal for two of these mutations differed between the complex environments and two simple environments in which we additionally manipulated opportunity for mate choice (via forced monogamy). We expected that if natural and sexual selection were aligned, we would see an increase in purging rate as accessibility to resources decreased and that the purging rate overall would be greater when sexual selection was allowed to act in the form of mate choice than when it was removed.

## Methods

### Environmental Manipulations

Images of the environmental treatments and an illustration of general set up are provided in Figure 1. Three environments were created in order to test the effects of desirable resource availability on the removal of deleterious mutations from populations. Within each environment there were both “high quality” resources of a yeast-rich food (see Table S1) and a 15% dilution (in water/carrageenan) of this food as a “low quality” resource. High quality food was determined based on previously published nutritional geometry studies (Lee et al. 2008; Jensen et al. 2015), that maximized female fecundity. Based on previous studies, the intent of these high quality food resources was to entice females to use these patches for oviposition and potentially lead males to defend these resources to maximize their own mating success. The diluted medium provided resource patches, such that individuals are not competing for survival *per se*, but for the desirable resources that females may prefer to maximize their fecundity. For each replicate environment described below, mesh BugDorm4M1515 cage (15cm^3^) were used. The “non-territory” treatment environment (NT) consisted of a single *Drosophila* culture bottle (177ml), with a surface area of 30.25cm^2^ (55mm × 55mm base) containing ~50ml of high quality food with the addition of four drops of a yeast-paste and orange juice mixture on top (to attract females (Dweck et al. 2013)), as well as a bottle only with 50ml low quality food. These represent “typical” *Drosophila* lab environments where apparent scramble competition is commonly observed (Spieth 1974), although subtle interference competition may be occurring as well (Baxter et al. 2018). The “unconstrained territory” spatial treatment (UCT) consisted of eight open vials (height of 50mm, 25mm diameter, 4.9 cm^2^ surface area) each filled with ~5 ml of high quality food with a single drop of yeast-paste/orange juice mixture on top and a single bottle with the low quality food. Finally, the “spatially constrained territory” treatment (SCT) had the same set-up as the UCT treatment except each vial had a 3D printed cap (22mm diameter, 25mm height, 4mm opening, see Supplemental Fig 1) to further restrict ease of access to high quality food patches. These 3D printed caps were designed and tested with several specific features in mind. First, that it was relatively difficult to gain access, but would be relatively easy (given positive photo-taxis and negative geo-taxis in *Drosophila* (Markow and Merriam 1977)) for an interloper to be chased out. Second, that the aperture was of sufficient size that two large *D. melanogaster* individuals could pass one another, but one individual could still harass or chase the other in this space. Finally, the cap was designed so that if an individual did display territorial behaviours, it had multiple places to survey or defend (food surface, inner aperture, and outside top of aperture). Pipe cleaners were wrapped around the tops of bottles and vials to provide additional perching substrate for individuals.

**Figure 1:**
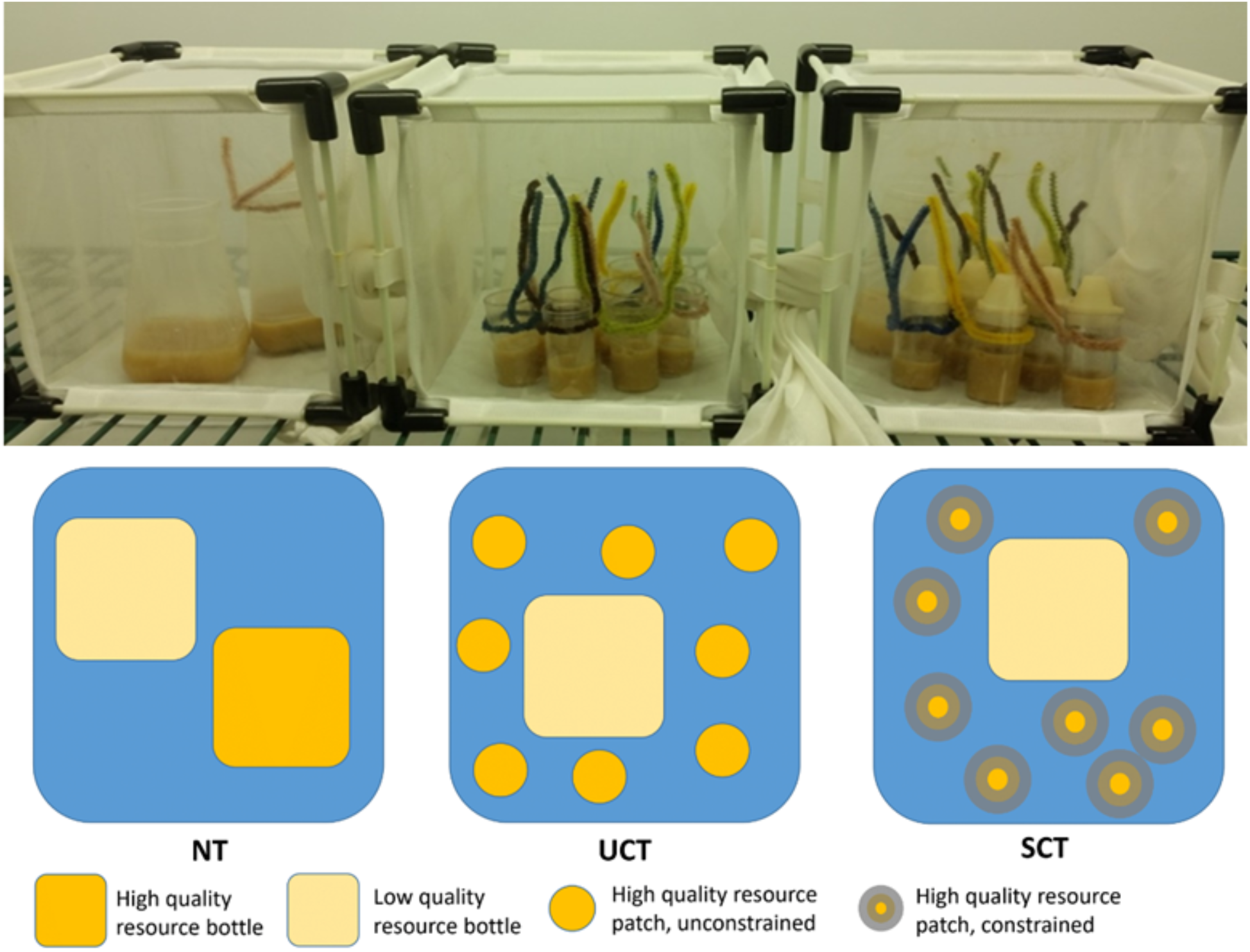
Top: Environmental treatment set up for non-territory (NT - left), unconstrained territory (UCT - middle), and spatially constrained territories (SCT - right). Bottom: Overhead schematic of environmental treatment set up.

### Populations

To examine deleterious allele purging rates, six mutations with known morphological defects were used across each of the three spatial treatments. Each allele was picked because of previous work examining the effects of selection on them in the context of either spatial manipulations or varying degrees of sexual selection (Arbuthnott and Rundle 2012; Colpitts et al. 2017). Three of these mutations are autosomal (*brown*^*1*^, *vestigial*^*1*^, and *plexus*^*1*^) and three are X-linked (*white*^*1*^, *yellow*^*1*^, and *forked*^*1*^). The mutations *plexus*^*1*^, *white*^*1*^, *yellow*^*1*^, and *forked*^*1*^ were obtained from Bloomington stock center while *brown*^*1*^ and *vestigial*^*1*^ were obtained from stocks kept in the lab. These alleles were chosen for their wide array of phenotypic effects with two influencing eye colour (*white*^*1*^ and *brown*^*1*^), two influencing wing morphology (*plexus*^*1*^ and *vestigial*^*1*^), one affecting body colour and behaviour (*yellow*^*1*^) and one affecting bristle morphology (*forked*^*1*^).

To create experimental populations, individuals were backcrossed into a large outbred domesticated lab population (census size of 1200-1600 individuals) originally collected from Fenn Valley Winery (FVW), Michigan (GPS co-ordinates: 42.578919, −86.144936) in 2010. This population was chosen to potentially minimize confounding effects of lab adaptation in this experiment (Harshman and Hoffmann 2000), i.e. it is expected that this population has already had considerable opportunity to adapt to our lab environment (~180 generations prior to initiation of this experiment). To generate experimental populations, the following procedure was used. For autosomal mutations, mutant female virgins were crossed with FVW males. F1 was then crossed to each other and mutant homozygote females were collected. For X-linked mutations, mutant males were crossed with wildtype females. The heterozygous females from this cross were then crossed back to wildtype males, the mutant offspring from this cross were then collected and the process was repeated. For each mutation, backcrossing was conducted for five generations and on the final generation, offspring from the final cross were mated together to create mutant males and females. Fifty pairs were used to generate each cross.

### Purging Rates Across Environments

For each mutation, nine replicate populations were created and three of each randomly assigned to one of the three environmental treatments. Initial populations consisted of 100 males and 100 females with starting allele frequencies of 0.7 for their respective mutation. Populations were maintained at 12L:12D cycles at 21°C with 60% relative humidity in a Conviron walk in chamber (CMP6050). Each generation, adults were placed into their respective treatments and allowed to mate and lay eggs for three days. After the three day period, adults were removed from the environments and discarded. Eggs were allowed to develop for 11 days, after which the next generation of adults was collected by bringing the adults to the cold room kept at 4°C and gently knocking them into vials. After this initial collection, 100 males and 100 females from each replicate were phenotyped under light CO2 and placed into their respective environments with fresh food. This cycle was repeated for 10 generations.

Due to a laboratory bacterial infection in one replicate of the *brown*^*1*^ population for the NT treatment, this replicate was discarded after generation 5. A fourth replicate was created with the same starting allele frequencies (0.7) in order to account for the missing data. This replicate was therefore five generations behind the rest of the experiment and was continued for 10 generations.

In order to get an estimate of allele frequencies for autosomal mutations during this experiment, monogamous pairings of phenotypically wildtype females and mutant males were conducted at generations 3 and 6, for *brown*^*1*^ and *plexus*^*1*^ populations and at generations 3 and 8 for *vestigial*^*1*^ populations. After the collection of adults for the next generation, for each population 50 virgin females were phenotyped over light CO2. Of the 50 females, those that lacked the mutation (i.e. could be homozygous or heterozygous for the wild type alleles) were placed singularly into vials with a mutant male. Offspring were analyzed from these vials over 3 days after emergence. If a vial contained only wildtype offspring, the female parent was scored as homozygous for lacking the mutation, if the vial contained a mixture of wildtype and mutant offspring, the female parent was scored as heterozygous for the mutation. For the X-linked mutations, allele frequencies were estimated from the frequency of the mutation in males.

### Purging Rates with Effects of Mate Choice

To determine the effects of sexual selection on purging rates, we re-ran the experiment using *white*^*1*^ and *vestigial*^*1*^ with the addition of two new treatments. The first treatment, deemed “vial no choice” (VNC), consisted of randomly assigning 100 individual pairs into vials to mate (i.e forced monogamy). The second treatment, “vial choice” (VC), consisted of randomly assigning 100 male and female adults into vials of 5 mixed sex pairs. After three days of mating for each treatment, males were removed and females were placed into environments similar to the NT treatment. After three days the females were removed and eggs were allowed to develop for 9-10 days. Emerging female virgins and adult males were collected similar to above and the process was repeated. NT, UCT, and SCT treatments were conducted the same as above except females were collected as virgins and males and females were held separately for three days after collection in order to align with the experimental schedule of the VNC and VC treatments. This experiment was conducted for only four generations as it was disrupted by a lab shutdown brought about by the covid-19 pandemic. One replicate of the SCT *vestigial*^*1*^ treatment did not have any surviving adults at generation four.

### Statistical Analysis

The rate of mutant allele loss in each population over multiple generations for each component of the experiment was analyzed by fitting generalized linear mixed effect models with binomial distribution (i.e. a logistic mixed model). Since each allele was started at a known frequency, and the intercept was known, models were fit without estimating a global intercept (but included offsets). Main effect for allele or treatment were also not included (as all treatments started with the same frequency for a given allele). Fixed effects included in the model were thus generation X mutation type, generation X treatment, and generation X mutant type X treatment. Random slopes for generation was included across replicate lineages, and the intercept was offset to 0.7 for allele frequency (or 0.49 for autosomal and 0.595 for sex-linked mutations when modelling mutant genotypic frequencies). Fixed effects were further examined for significance with a two way ANOVA (type II Wald χ^2^ test) and treatment contrasts averaged over mutant type were examined by comparing estimated marginal means within each model.

For analyzing purging rates across environmental treatments, models were generated with and without the third SCT replicate for the *forked*^1^ mutation due to this replicate having mutant allele frequencies approaching fixation consistently throughout the experiment (Figure S2, Table S2 and Table S3). Results presented exclude this replicate unless otherwise indicated.

Selection coefficients for each mutation were estimated using the allele frequency data. Selection coefficient per generation was calculated as s = 1 – (q’/q), and these estimates were then averaged across generation and replicate for each mutant type.

All statistical analyses were performed in R v.3.5.2 (R Core Team 2018) using glmer() (lme4 package v1.1-21 (Bates et al. 2015)), Anova() (car package v3.0-2 (Fox and Weisberg 2011)), and emtrends() (emmeans package v.3.1.1 (Lenth 2019)). All plots were generated with ggplot2 v.3.1.1 (Wickham 2016).

## Results

As expected, average allele frequency declined over generations for all six mutations types, indicating these alleles to be deleterious (Fig 2). We observed substantial differences in rates of purging (as assessed by genotypic frequencies) based on the identity of the mutation. ANOVA shows significant effects for all interactions of generation with mutant type and treatment, however significant effects may be restricted to certain mutation types as contrast estimates between treatments among all mutant types are non-significant (Table 1 and Table 2). Across the six mutation types, there was no consistent overall pattern in purging rate between the NT, UCT, and SCT environmental treatments. Similar results are shown when analyzing males and females separately. When examining estimated allele frequencies, only the interactions between generation and mutant type, and generation and treatment are significant (Fig 3, Table 1). However, treatment contrasts are still not significantly different from one another (Table 2). Overall trends of significance from ANOVA and treatment contrasts are the same when including the third SCT replicate for the *forked*^1^ mutation. Estimated selection coefficients are of differing strengths for each mutant type, however these estimations also indicate no consistent pattern in strength of selection of treatment types across mutations (Fig 4). Overall the results suggest that while there are effects of the three spatial treatments on rates of purging (Fig S3), they are relatively modest in comparison to the effects of individual mutants and their interactions with the spatial treatment.

**Figure 2:**
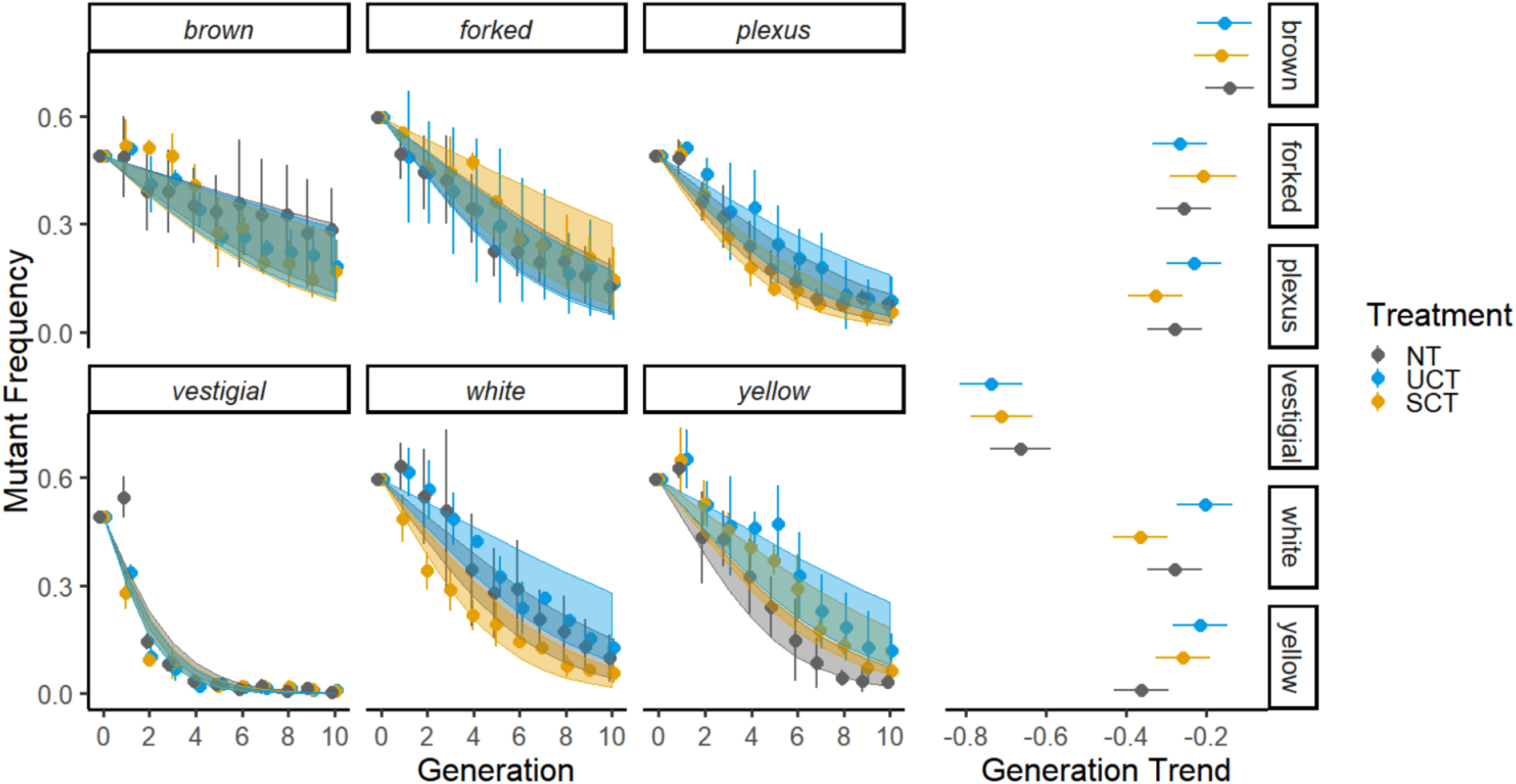
Left: Purging rates across the three environmental treatments for each mutant. Data points and error bars represent mean mutant frequency and standard deviation across the three replicates. Confidence bands represent 95% confidence intervals for our generalized linear mixed model. Right: Treatment contrasts for each mutant type based on model estimates. Error bars represent 95% confidence intervals.

**Table 1:**
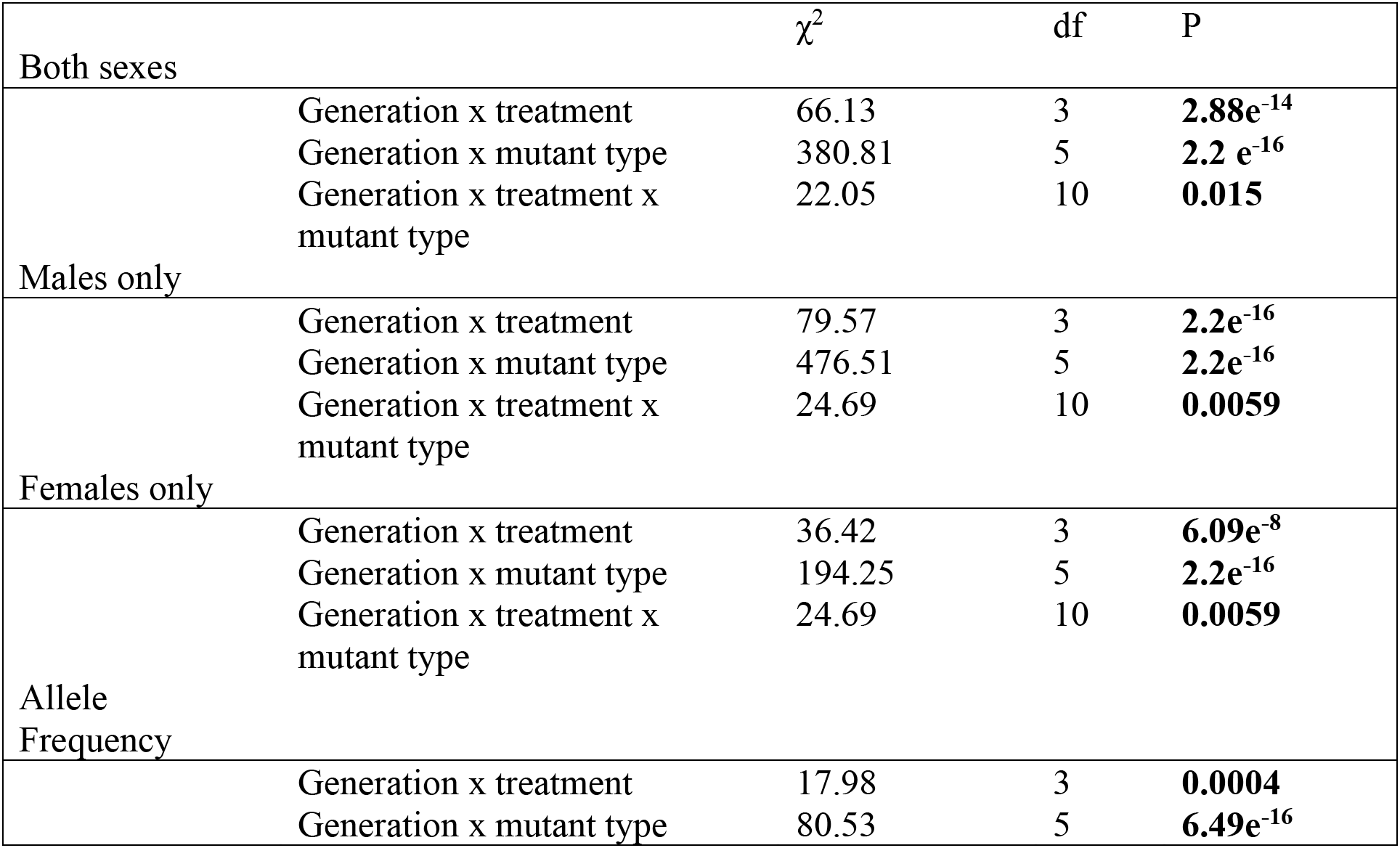
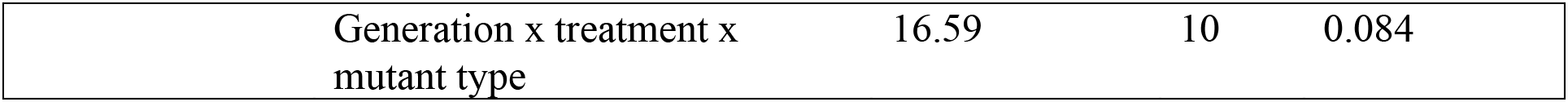
ANOVA outputs for fixed effects of four general linear mixed models produced from the six mutant types across the three treatment types.

**Table 2:**
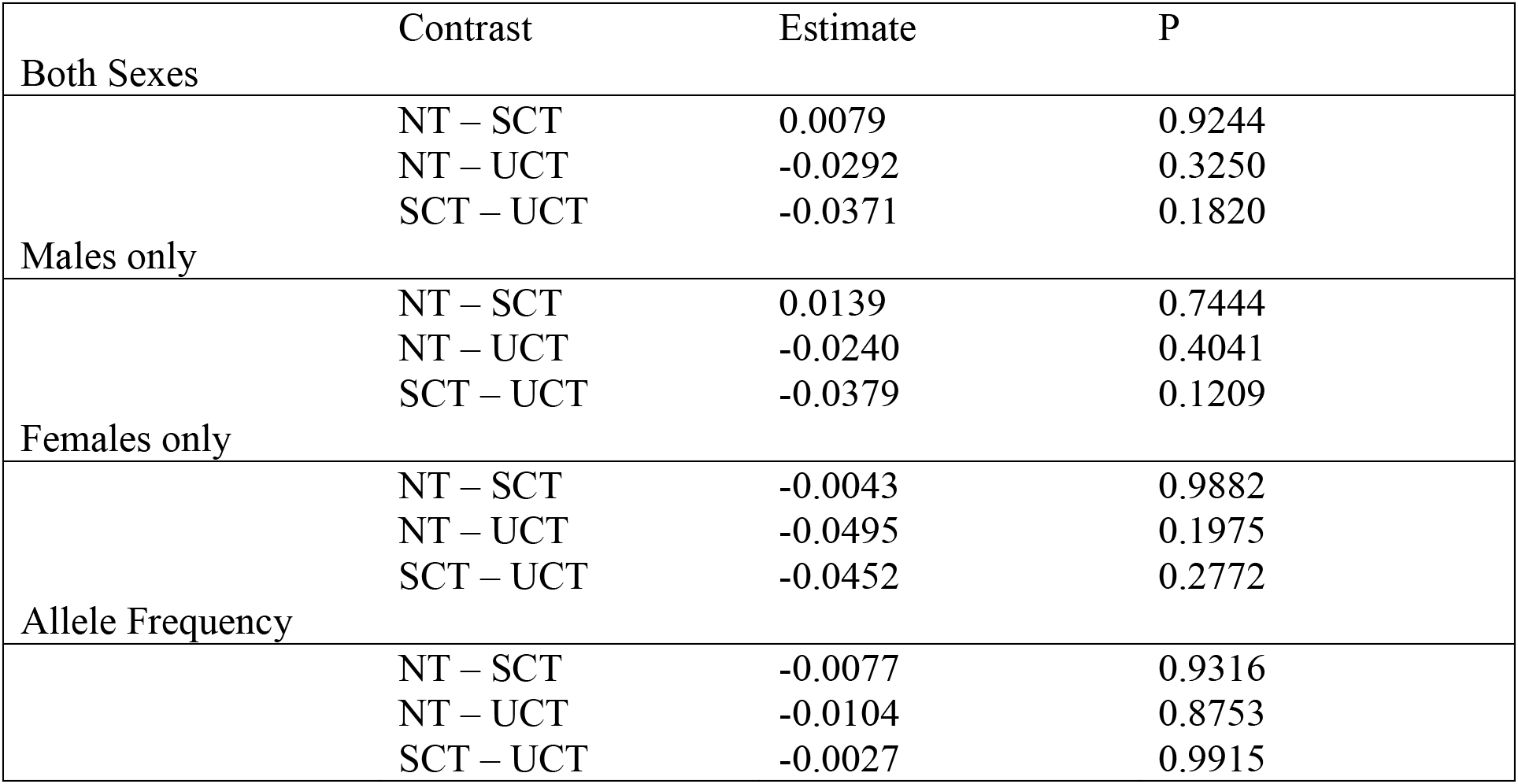
Estimates and significance of treatment contrasts among the six mutation types for four general linear mixed models

**Figure 3:**
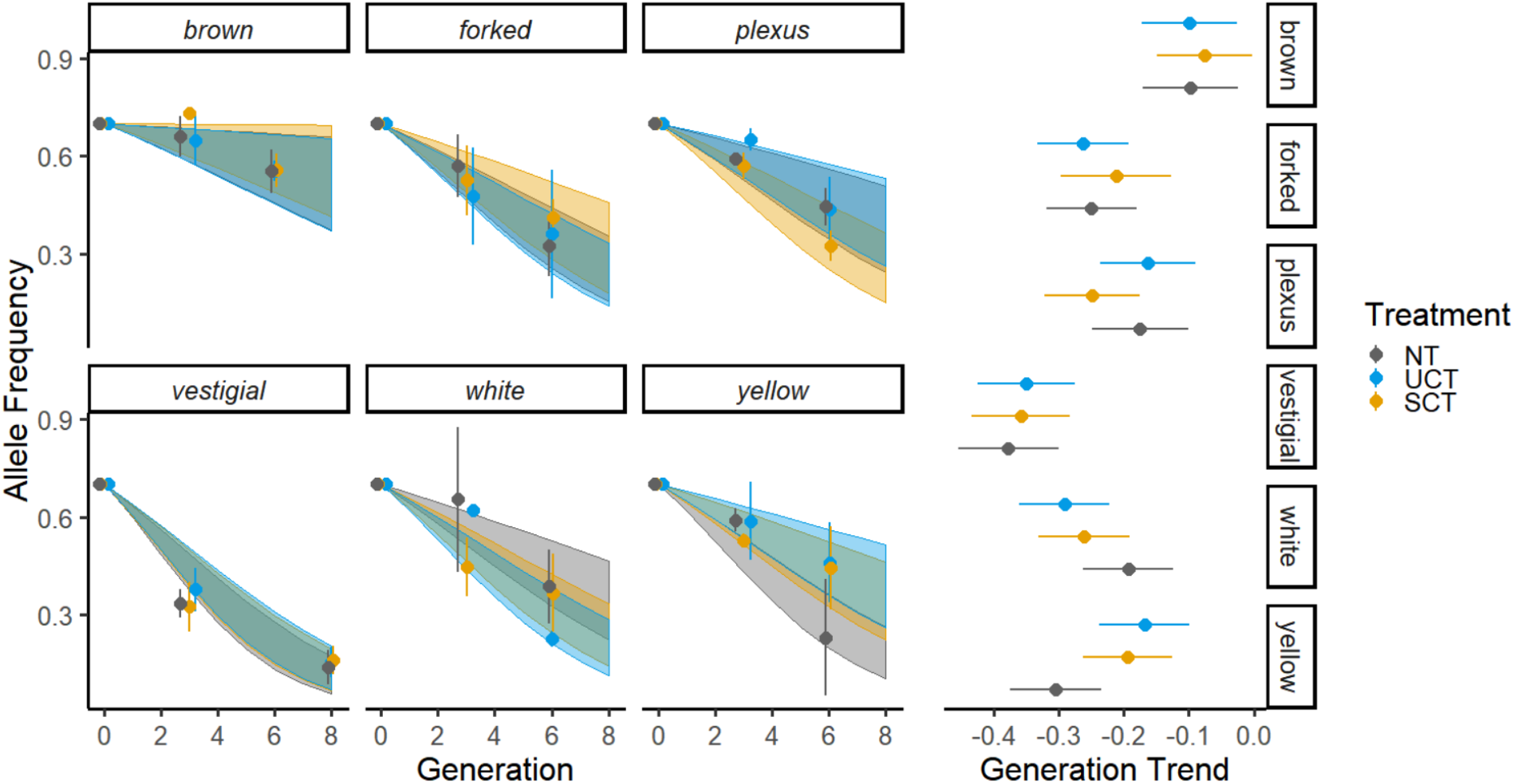
Left: Purging rates across the three environmental treatments for each mutant. Data points and error bars represent mean allele frequency and standard deviation across the three replicates. Confidence bands represent 95% confidence intervals for our generalized linear mixed model. Right: Treatment contrasts for each mutant type based on model estimates. Error bars represent 95% confidence intervals.

**Figure 4:**
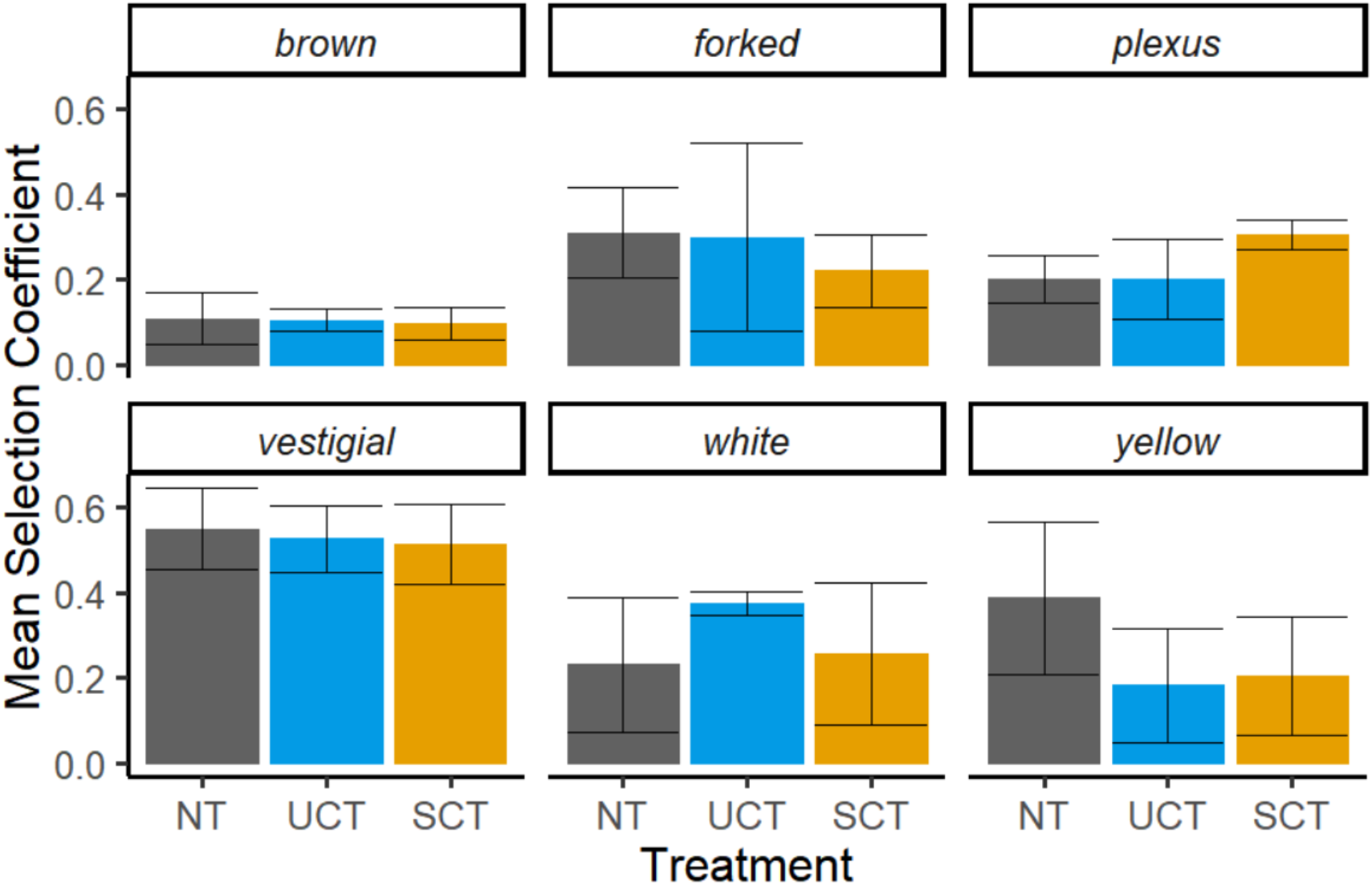
Mean selection coefficients for each mutant type across the three environmental treatments. Estimates were created from allele frequency data, error bars represent 95% confidence intervals.

In the second experiment, we replicated the above experiment with two alleles and added additional treatments with explicit manipulations of sexual selection. The addition of sexual selection for both *white*^*1*^ and *vestigial*^*1*^ mutant populations increased purging rates (Fig 5, Table 3). While the forced monogamy treatment (VNC) treatment showed the slowest purging rate for both mutations, between the treatments that include sexual selection there is no consistent pattern in purging rate by treatment across the two mutant types. The ANOVA shows significant effects of the interaction between generation and mutant type, and generation and treatment but not for the interaction between all three fixed effects. Treatment contrasts show that the VNC (vial no choice) treatment (i.e. forced monogamy) is significantly different from the other treatments but VC, NT, UCT, and SCT are not significantly different from each other. When analyzing the sexes separately, only the interactions between generation and treatment, and generation and mutant type were significant for males whereas the interactions between generation and treatment, and generation, treatment and mutant type were significant for females. Treatment contrasts were similar between male and female models with only the VNC treatment showing a significant difference from other treatment types when looking across all mutation types (Table 4).

**Figure 5:**
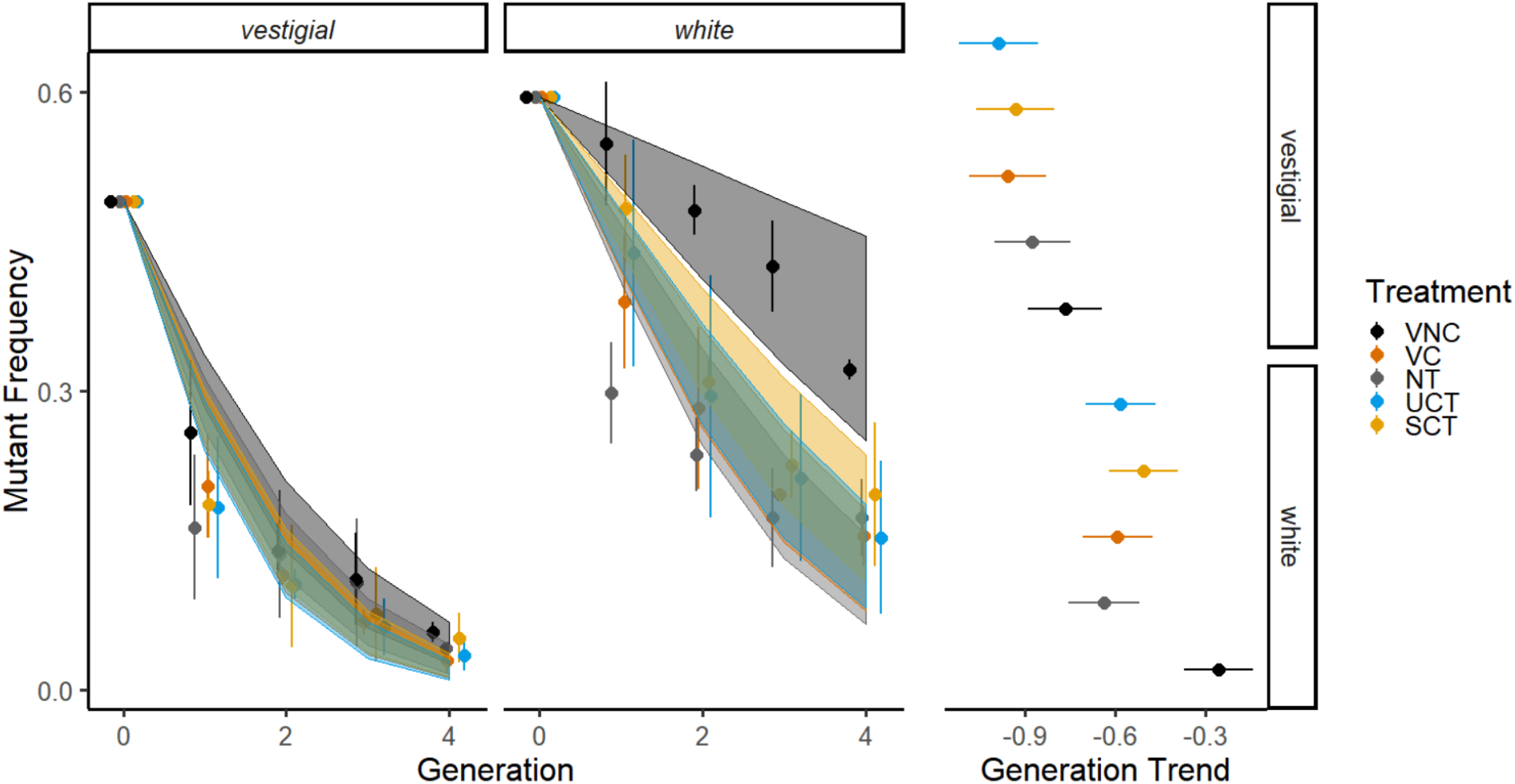
Left: Purging rates across the five environmental treatments for each mutant. Data points and error bars represent mean mutant frequency and standard deviation across the three replicates. Confidence bands represent 95% confidence intervals for our generalized linear mixed model. Right: Treatment contrasts for each mutant type based on model estimates. Error bars represent 95% confidence intervals

**Table 3:** ANOVA outputs for fixed effects of three general linear mixed models produced from the two mutant types across the five treatment types.

| | | $\chi^2$ | df | P |
| --- | --- | --- | --- | --- |
| Both sexes |  |  |  |  |
|  | Generation x treatment | 973.56 | 5 | **2e<sup>-16</sup>** |
|  | Generation x mutant type | 97.78 | 1 | **2e<sup>-16</sup>** |
|  | Generation x treatment x mutant type | 5.21 | 4 | 0.27 |
| Males only |  |  |  |  |
|  | Generation x treatment | 1272.95 | 5 | **2e<sup>-16</sup>** |
|  | Generation x mutant type | 460.20 | 1 | **2e<sup>-16</sup>** |
|  | Generation x treatment x mutant type | 8.85 | 4 | 0.065 |
| Females only |  |  |  |  |
|  | Generation x treatment | 431.31 | 5 | **2e<sup>-16</sup>** |
|  | Generation x mutant type | 1.03 | 1 | 0.311 |
|  | Generation x treatment x mutant type | 12.39 | 4 | **0.015** |

**Table 4:** Estimates and significance of treatment contrasts among *white*^*1*^ and *vestigial*^*1*^ mutations for three general linear mixed models

|  | Contrast | Estimate | P |
| --- | --- | --- | --- |
| **Both Sexes** |  |  |  |
|  | VNC – NT | 0.2451 | **0.0007** |
|  | VNC – VC | 0.2633 | **0.0002** |
|  | VNC – SCT | 0.2074 | **0.0072** |
|  | VNC – UCT | 0.2733 | **0.0001** |
|  | NT – VC | 0.0183 | 0.9984 |
|  | NT – SCT | -0.0377 | 0.9750 |
|  | NT – UCT | 0.0282 | 0.9917 |
|  | VC – SCT | -0.0560 | 0.9014 |
|  | VC – UCT | 0.0099 | 0.9999 |
|  | SCT – UCT | 0.0659 | 0.8355 |
| **Males Only** |  |  |  |
|  | VNC – NT | 0.2142 | **0.0010** |
|  | VNC – VC | 0.3389 | **<0.0001** |
|  | VNC – SCT | 0.1712 | **0.0197** |
|  | VNC – UCT | 0.2844 | **<0.0001** |
|  | NT – VC | 0.1246 | 0.2709 |
|  | NT – SCT | -0.0430 | 0.9475 |
|  | NT – UCT | 0.0701 | 0.7747 |
|  | VC – SCT | -0.1676 | 0.0639 |
|  | VC – UCT | -0.0545 | 0.9214 |
|  | SCT – UCT | 0.1131 | 0.3507 |
| **Females Only** |  |  |  |
|  | VNC – NT | 0.3555 | **0.0001** |
|  | VNC – VC | 0.3392 | **0.0003** |
|  | VNC – SCT | 0.3007 | **0.0021** |
|  | VNC – UCT | 0.3501 | **0.0002** |
|  | NT – VC | -0.0163 | 0.9997 |
|  | NT – SCT | -0.0548 | 0.9664 |
|  | NT – UCT | -0.0055 | 1.0000 |
|  | VC – SCT | -0.0385 | 0.9908 |
|  | VC – UCT | 0.0108 | 0.9999 |
|  | SCT – UCT | 0.0493 | 0.9769 |

## Discussion

Spatial heterogeneity in the environment can alter many aspects of an organisms’ phenotype including mating strategy which in turn influences how selection acts on a population including the degree to which allelic effects may be concordant or antagonistic across fitness components. Analyzing the directions and magnitudes of the components of natural selection has been investigated in many contexts, however many empirical studies teasing apart these elements in varying environments fail to recognise the influence of mating strategies. We created populations with known mutation frequencies and allowed them to evolve in environments differing in spatial constrains for resource accessibility to determine how environmental complexity influences the removal of deleterious mutations. We found environmental complexity did influence purging rates, but these rates depended greatly on mutation type. We reanalyzed the purging rates of two of these mutations in the same environments but also including treatments allowing different opportunities for mate choice within a more “simple” environment. Again, we found that purging rates between treatments varied with mutation type, but for both mutations a lack of mate choice (forced monogamy) decreased purging rates.

For each of the six mutations, we expected that with increased variance in resource accessibility there would be an increase in purging rate and therefore the highest purging rate would be seen in the SCT treatment, with the lowest being in the NT treatment. This prediction rested on several assumptions including that natural and sexual selection are aligned, high quality food patches in the SCT treatment would initiate territorial behaviour within males, and males of the highest quality would be able to hold and defend these food patches with the most success, leading to the most mates. While the SCT treatment showed the highest purging rate among treatment types for *plexus*^*1*^ populations, this pattern does not hold for other mutant types. This discrepancy between our predictions and the data could be due to inaccurate assumptions or other unknown factors. Despite evidence that *Drosophila melanogaster* among other *Drosophila* species can show context dependent territoriality (Hoffmann 1987; Hoffmann and Cacoyianni 1990), considerable uncertainty exists in the extent of what factors influence it and how it ultimately influences the fitness of an individual. It should also be noted that evolutionary stable strategy theories predict that a behavioural strategy will only be adopted by an individual or population if it is advantageous (Maynard Smith 1974). While our environments were designed based on theory that would suggest our assumptions provide the most advantageous strategy (Emlen and Oring 1977; Emlen 2014), this cannot be known without further empirical testing and observation and other strategies may have been implemented that cause the discrepancy between our expectations and results.

The lack of consistency between mutant alleles and the difference between treatments could be due to populations not using the environments as predicted. The NT environment was designed to resemble environments that promote scramble competition in *Drosophila*, with UCT having characteristics that promote territorial behaviours. The SCT environment was designed to provide greater opportunity for one-on-one contests to occur between individuals due to limited entry to the desirable resource. Individuals in the UCT and SCT environments were provided 25mm diameter high quality food patches with potential densities of 12 males per high quality food patch (if the 100 males within each environment were equally distributed across patches). While these conditions have been shown to increase the rate of territorial behaviour and the success of those males that defend territories (Hoffmann and Cacoyianni 1990), these results were found over short-term experiments (up to 6 hours) and these behaviours may not persist in *D. melanogaster* populations over longer time periods like the three days we allowed in our experiment. Although not observed, other unexpected uses of the environments such as the majority of copulations occurring outside of food patches, and skewed patch use could have caused the disparity between our predictions and results. Also, the addition of cap in the SCT treatment was expected to aid males in further defending their resource patches. However due to the novelty of these environments, the behaviours these environments were meant to encourage may not have been used or had the opportunity to evolve. If the behaviours did evolve but at a point in the experiment where the allele frequencies for the mutations were low, genetic drift could have masked their effects.

Although our results do not show any consistent pattern of purging rate across treatment types between mutant types, inconsistent results are common to many purging experiments. Many studies that analyze multiple mutations find that each mutation acts differently to experimental treatments not only in magnitude but also direction and thus mainly focus on the overall patterns among mutation types (Sharp and Agrawal 2008; MacLellan et al. 2009; Arbuthnott and Rundle 2012; Clark et al. 2012; Maclellan et al. 2012; Colpitts et al. 2017; Singh et al. 2017). These differences are also reflected in our calculated selection coefficients, where higher selection coefficients lead to faster purging rates but the environmental treatment that has the highest selection coefficient changes depending on mutation type. Differences between how these mutant individuals interact within their environment can likely explain these variances. For example, the mutant *vestigial* has a wing phenotype that influences both its movement and courtship signalling (Pezzoli et al. 1986) putting it at a greater disadvantage compared to wild-type individuals in the same population, which is likely why it has the most drastic purging rate across environmental treatments among all the mutations analyzed in this study. Further investigation into the behaviours of these mutant types may give an indication as to why these results differ between mutant types.

While we wanted to explore how resource accessibility and environmental complexity influence populations through purging rates, we also wanted to evaluate how these compared to the purging rates of populations that lacked sexual selection and populations that had simple mating environments. As expected, the addition of sexual selection increased the purging rate for both mutations tested. However, there was no difference between the simple and relatively complex environments in purging rate for either mutation. This contradicts previous work of Colpitts et al. (2017) where polygamous populations of mutant *white*^*1*^ *D. melanogaster* showed increased purging rates in complex environments. While the overall ideas between our experiments are similar, key differences in experimental design could explain these differences. Firstly, due to the alignment of the experimental schedule, virgins from the VNC and VC treatments were able to mate more quickly than the virgins in the NT, UCT, and SCT treatments that were initially held separately before mating. This difference in waiting times to mate could have caused virgins from the NT, UCT, and SCT treatments to be more receptive to potential mates (Pavković-Lučić and Kekić 2009). This could also explain why we see differences in the overall trends between the NT, UCT, and SCT environments compared to our initial experiment. Secondly, our experiment had a much shorter mating period (3 days versus 6) and all eggs laid during this time period were kept to potentially contribute to the next generation for the NT, UCT, and SCT treatments, but not for the VNC and VC treatments. This could potentially lead to lower quality offspring from early matings with lower quality males being kept within the experiment, decreasing the purging rates within the complex mating treatments.

Overall our study adds to the recently growing body of literature considering “environmental complexity” while breaking down “complexity” further to accommodate for changes in mating strategy by environment.

## Acknowledgements

We thank Dr. Tony Frankino and Christine Sikes for their assistance with the design and development of 3D-printed caps used in this study. Funding for this research was provided by the Natural Sciences and Engineering Research Council (NSERC) of Canada and McMaster University to ID.

## Supplementary

**Figure S1:**
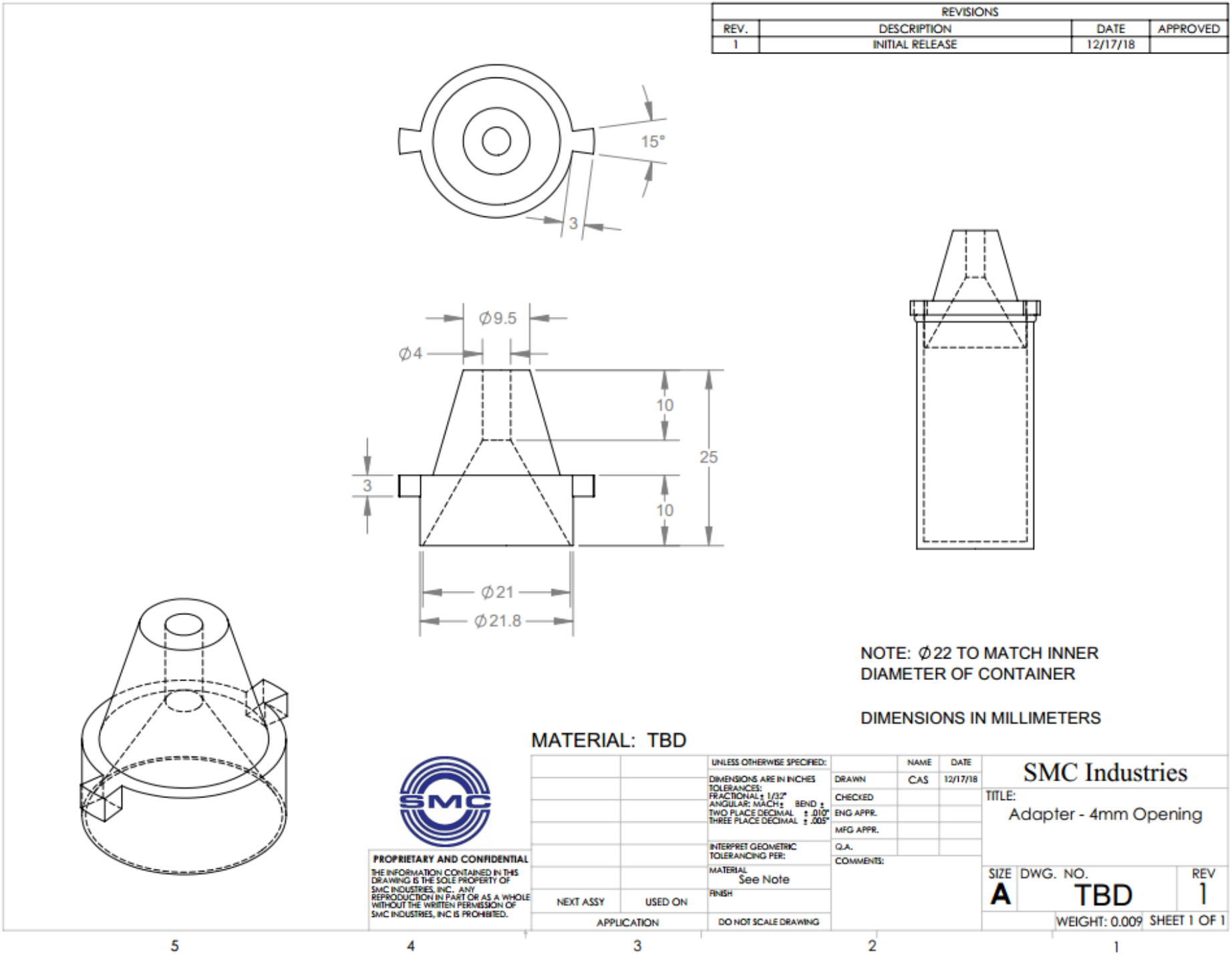
Schematic for 3D-printed cap design. Caps were created using filament material.

**Figure S2:**
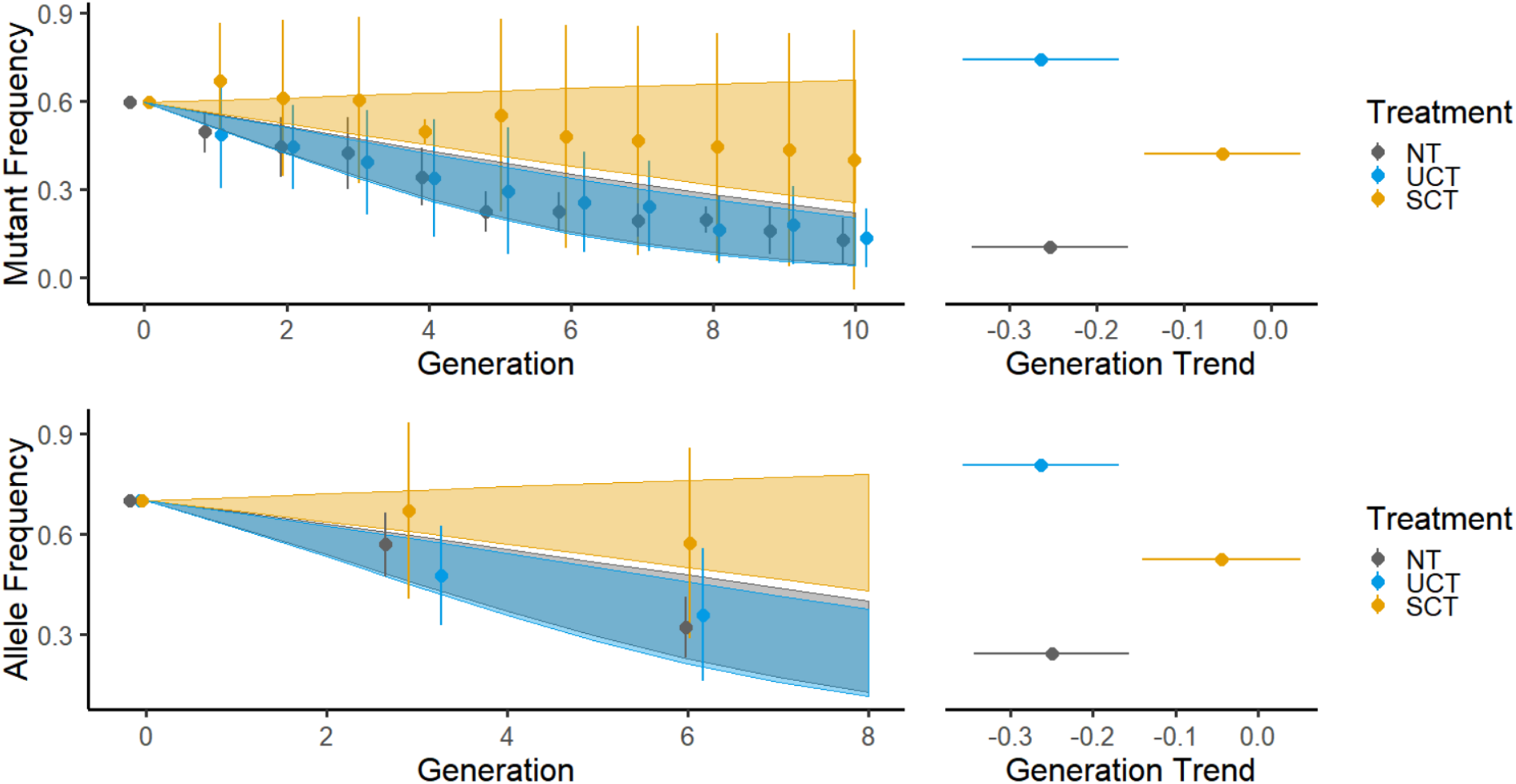
Right: Purging rates for *forked*^*1*^ mutant while including all replicates. Data points and error bars represent average mutant frequency or allele frequency and the standard deviation across all replicates. Confidence bands represent 95% confidence intervals for our generalized linear mixed model. Left: Treatment contrasts for *forked*^*1*^ mutant while including all replicates based on model estimates. Error bars represent 95% confidence intervals

**Figure S3:**
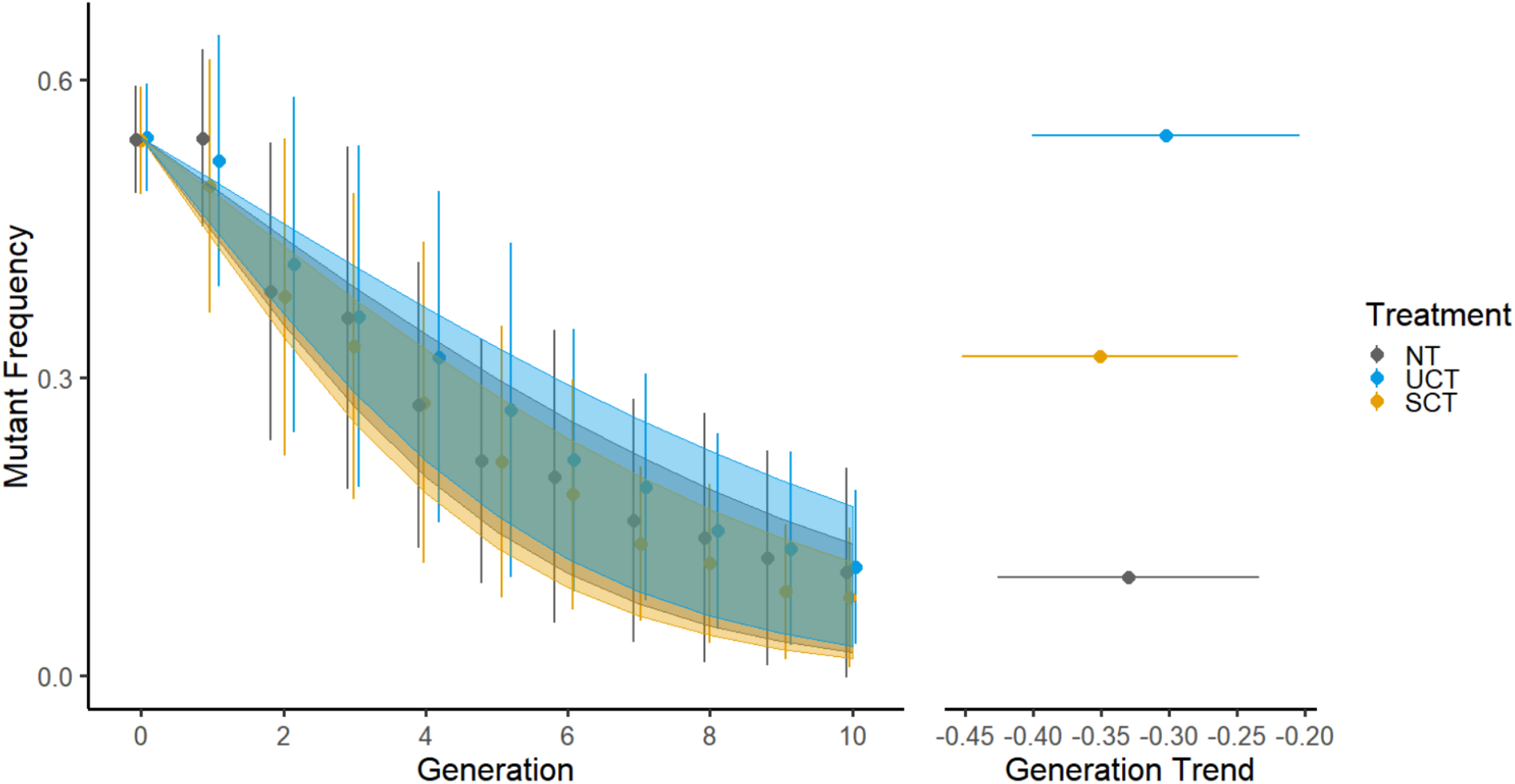
Left: Purging rates across the three environmental treatments averaging across mutant types. Data points and error bars represent mean mutant frequency and standard deviation across the three replicates of each mutant type. Confidence bands represent 95% confidence intervals for our generalized linear mixed model treating mutant type as a random effect. Right: Treatment contrasts across mutant types based on model estimates. Error bars represent 95% confidence intervals.

**Table S1:** Recipe for high quality food and nutritional contents

| Ingredient | Mass (g) | Volume (ml) | Carbohydrates (g) | Protein (g) |
| --- | --- | --- | --- | --- |
| Water | 4250 | 4250 |  |  |
| Black strap molasses | 40 | 28 | 24.40 | 1.60 |
| Fancy table molasses | 40 | 28 | 30 | 0.40 |
| Cornmeal | 70 | 106.06 | 56.70 | 4.90 |
| Carrageenan | 27 | 19.71 |  |  |
| Yeast | 255 | 447.37 | 84.15 | 127.50 |
| Sucrose | 75 | 46.88 | 75 | 0 |
| Propionic acid | 11.85 | 12 |  |  |
| Methylparaben | 2.50 | 1.81 |  |  |
| Ethanol | 19.61 | 25 |  |  |
| TOTAL | 4790.97 | 4964.85 | 270.25 | 134.4 |

**Table S2:** ANOVA outputs for fixed effects of four general linear mixed models produced from the six mutant types across the three treatment types including all replicates.

| | | $\chi^2$ | df | P |
| --- | --- | --- | --- | --- |
| Both Sexes |  |  |  |  |
|  | Generation x treatment | 37.99 | 3 | **2.84e<sup>-8</sup>** |
|  | Generation x mutant type | 253.39 | 5 | **2e<sup>-16</sup>** |
|  | Generation x treatment x mutant type | 26.72 | 10 | **0.003** |
| Males Only |  |  |  |  |
|  | Generation x treatment | 36.21 | 3 | **6.77e<sup>-8</sup>** |
|  | Generation x mutant type | 295.41 | 5 | **2e<sup>-16</sup>** |
|  | Generation x treatment x mutant type | 26.24 | 10 | **0.003** |
| Females Only |  |  |  |  |
|  | Generation x treatment | 25.29 | 3 | **1.34e<sup>-5</sup>** |
|  | Generation x mutant type | 142.61 | 5 | **2e<sup>-16</sup>** |
|  | Generation x treatment x mutant type | 30.93 | 10 | **0.0006** |
| Allele Frequency |  |  |  |  |
|  | Generation x treatment | 11.81 | 3 | **0.008** |
|  | Generation x mutant type | 47.52 | 5 | **4.46e<sup>-9</sup>** |
|  | Generation x treatment x mutant type | 19.84 | 10 | **0.032** |

**Table S3:** Estimates and significance of treatment contrasts among the six mutation types for four general linear mixed models including all replicates

|  | Contrast | Estimate | P |
| --- | --- | --- | --- |
| Both Sexes |  |  |  |
|  | NT – SCT | -0.0181 | 0.7741 |
|  | NT – UCT | -0.0299 | 0.4997 |
|  | SCT – UCT | -0.0118 | 0.8998 |
| Males Only |  |  |  |
|  | NT – SCT | -0.0150 | 0.8422 |
|  | NT – UCT | -0.0258 | 0.6020 |
|  | SCT – UCT | -0.0108 | 0.9162 |
| Females Only |  |  |  |
|  | NT – SCT | -0.0328 | 0.6024 |
|  | NT – UCT | -0.0504 | 0.3034 |
|  | SCT – UCT | -0.0176 | 0.8664 |
| Allele Frequency |  |  |  |
|  | NT – SCT | -0.0358 | 0.4162 |
|  | NT – UCT | -0.0104 | 0.9278 |
|  | SCT – UCT | 0.0254 | 0.6431 |

